# Quantitative ethology of schistosome miracidia characterizes a conserved snail peptide that inhibits penetration

**DOI:** 10.1101/2025.07.02.662618

**Authors:** Rachel V. Horejsi, Chase N. Nelson, Avery De Ruyter, Helen Gensch, Saige Maasz-Seawright, Carly Weber, Sophie Willett, Sonja A. Olson, Nicolas J. Wheeler

**Affiliations:** Department of Biology, University of Wisconsin-Eau Claire, Eau Claire, WI

## Abstract

Over 700 million people are at risk of contracting schistosomiasis due to regular exposure to freshwater sources where infected snails, the obligate intermediate hosts of schistosomes, are endemic. Although mass drug administration of praziquantel effectively controls the disease in most regions, achieving elimination will require reducing populations of infected snails that shed the human-infective larval stage. Considerable effort has focused on parasite development and immunological responses after snail penetration, but comparatively little is known about the molecular and behavioral host seeking events that precede it, primarily due to technical and physical constraints. To address this gap, we developed a custom imaging and computational system for tracking and screening schistosome miracidia, the snail-infective larval form that hatches from eggs. Our system employs an array of cameras without magnification and acrylic devices that maintain miracidia within the focal plane, create a field of view over 200,000 times the area of a single miracidium, and support the formation of stable chemical gradients. Using this platform, we perform quantitative ethology of miracidia at an unprecedented scale and extract features that drive the emergent chemoklinokinetic behavior in response to snail cues. We demonstrate that miracidia accumulate at the edge of a gradient of snail cues by increasing key chemoklinokinetic features upon leaving the region of a cue, corroborating previous reports. In contrast, miracidia do not exhibit these behaviors when the cue is uniform, demonstrating that they represent a specific sensory response rather than generic neuromuscular activity. We further find that a previously identified stimulatory snail peptide only partially recapitulates the full chemoklinokinetic profile, and homologues from closely related species elicit divergent behavioral outcomes. Notably, some of these snail peptides can mask a natural gradient and inhibit miracidia penetration of snails. This work establishes a scalable behavioral platform for probing parasite- snail interactions and identifies a peptide scaffold that potently blocks snail penetration.

## Introduction

Schistosomiasis is a highly prevalent neglected tropical parasitic disease that infects over 250 million people worldwide, with hundreds of millions more at risk. Low-income communities without adequate access to clean water and sanitation are disproportionately impacted (1,2). Sustained mass drug administration (MDA) of praziquantel (PZQ) has reduced schistosomiasis infection prevalence and intensity, but because of a lack of control efforts targeting the transmission cycle itself, reinfection and persistent hotspots are common (3,4). Schistosomes, the parasitic flatworms that cause schistosomiasis, require the progression through a snail intermediate host to become infective to humans, making the snail-infective stage a chokepoint that, when targeted for control, greatly increases the effectiveness of MDA and reduces prevalence even in the absence MDA (5). Though much effort has been invested in understanding schistosome development and snail immunological responses following penetration, comparatively little is known about the host-seeking behaviors and molecular events that prelude snail infection.

Miracidia, the free-living, aquatic larvae that hatch from eggs and infect snails, exhibit behavioral alterations in response to chemical signals that indicate a nearby snail host. Most ethological studies of miracidia behavior have relied on qualitative observations that highlight specific features of swimming alterations and accumulation tendencies (6–9). More recent studies have used video recording and image analysis software to quantitatively describe miracidia behavior (10–13). From this work, miracidia have been shown to perform chemoklinokinesis when they are exposed to snail cues – they increase the frequency and magnitude of their turns and the tortuosity of their swimming paths, leading to accumulation in areas of higher concentration (14). However, throughout the history of miracidia behavioral parasitology, many reports have been difficult to reproduce due to the variety of experimental approaches and limited rigorous quantitative analyses. For example, some evidence suggests that snail cues do not cause a chemoklinokinetic behavior when presented uniformly, while others have shown the opposite (9,15). Still other reports suggest that miracidia change their behavior only when moving up a concentration gradient rather than down it (16). Conflicting reports have led to the preference for assays that measure the emergent accumulation phenotype rather than the behaviors that drive this phenotype (17).

Likewise, little is certain about the specific snail cues that elicit the chemoklinokinetic response in nature, as there are many reports regarding potential bioactive snail molecules (18–21). Recently a snail-secreted peptide known as P12 was identified and was shown to elicit similar behavioral responses to those caused by snail mucus and snail-conditioned water, suggesting it may play a role in miracidia host-seeking (10). However, the molecular and ethological details of the role of P12 have not been studied.

Much of our current understanding of miracidia ethology has been a result of studies evaluating behavior for short time frames and within small fields of view. This limitation is due to physical and technical constraints that make relevant spatiotemporal imaging scales challenging to achieve.

Maintaining resolution and magnification within a large enough field of view to accurately capture behavioral profiles of miracidia when they encounter cues has proven challenging due to their small size (miracidia are <200 µm long) and their ability to swim in all three dimensions in an aquatic environment. Together, these factors make it difficult to keep miracidia in the focal plane during recording and ensure they are not hidden by shadows. Additionally, the establishment of stable chemical gradients in liquid is difficult to control and reliably reproduce. These limitations have resulted in ethological studies that have often relied on qualitative observations or quantitative measurements limited in scale.

We have developed an imaging platform that employs an array of high-resolution cameras and customizable acrylic arenas that expand the field of view to an unprecedented scale, maintain miracidia within the focal plane, and support stable and reproducible chemical gradients. Using this platform and an expanded quantitative behavioral feature set, we describe the behaviors of miracidia as they explore a large ethology arena with competing chemical cues. To complement the ethology arenas, we have designed a screening arena that allows for high-throughput behavioral screens of miracidia. The combination of these approaches allows us to sensitively describe the chemoklinokinetic features in response to cue gradients or chemical treatments, leading to the identification of molecules that can act on miracidia to block the penetration of snails.

## Methods

### Snail husbandry and parasite maintenance and harvesting

*Biomphalaria glabrata* (NMRI strain) was maintained in two separate aquaria: a 30-gallon main colony tank and a 20-gallon nursery for eggs and juvenile snails. Both tanks were filled with artificial pond water (APW) and maintained at 25°C with constant aeration. Snails were fed three times per week with one large leaf of organic romaine lettuce per tank and supplemented with a small amount of fish food (Tetramin). Egg masses were deposited on floating pieces of Styrofoam. Styrofoam pieces with high densities of viable egg sacs were transferred to the nursery tank for hatching. Juveniles were returned to the main tank once sufficiently grown.

Livers from *S. mansoni* (NMRI)-infected female Swiss outbred mice (The Jackson Laboratory) were harvested at the Biomedical Research Institute at 6 weeks post-infection and shipped overnight in perfusion fluid to UWEC. Miracidia were harvested as previously described (22,23). Miracidia density was calculated by counting five 10 µL aliquots stained with 1:1 Lugol’s iodine and diluted with fresh APW to a final density of 1 parasite/µL to use in ethology assays and variable final densities for high-throughput screening assays. Miracidia were directly transferred from the harvesting flask to a glass petri dish and individually pipetted for penetration experiments.

### InVision design

For high-resolution wide-field imaging, we designed the InVision (for <u>inv</u>ertebrate <u>vision</u>) by customizing a Kastl-HighRes unit controlled by the Motif software (Loopbio GmbH, Vienna, Austria) based on inspiration from similar recording devices for *C. elegans* (24). The InVision system includes four cameras (Basler acA5472-17um, Ahrensburg, Germany) and lenses (Fujifilm Fujinon CF50ZA-1S, Tokyo, Japan) affixed to T-slotted extrusion bars in an aluminum enclosure (S1 Fig). The enclosure is not environmentally controlled but has two exhaust fans to maintain ambient temperature. Arenas sit upon a laser-cut black acrylic stage with a 200 x 200 mm white, infrared (850 nm), or red (625 nm) LED panel (MBJ DBL-2020 series, Ahrensburg, Germany) below to generate a homogenous bright background by transillumination. The InVision also contains RGB LEDs on mounts to provide epi-illumination, but the functionality was not used for this project.

Cameras are paired to create two distinct fields of view (FoV) of 81×27 mm, with the FoV of paired cameras overlapping ∼10%. The 1” camera sensors at 200 mm working distance creates an image with a resolution of 126.5 px/mm, meaning a miracidium consists of an ellipsoid ∼25 pixels long and ∼8 pixels wide. The Motif software synchronizes camera capture while recording frame-wise environmental variables via Phidget sensors (humidity, temperature, and luminosity). Motif utilizes GPUs to compress videos in real time according to the HEVC (H.265) standard and stores them in a MP4 container. HEVC efficiency relies upon a homogenous background, and our arena design (see below) routinely allowed compression ratios up to 100:1 without sacrificing tracking of extremely small animals. Videos are stored on a network attached storage device (Synology, New Taipei City, Taiwan) and transferred to the UWEC BOSE high-performance computer for tracking analysis.

### Arena and agarose mold design and fabrication

Design files (PDF, STL, or CAD) for arenas and molds can be found in the associated GitHub repository (wheelerlab-uwec/miracidia-sensation-ms). All arena parts were fabricated from 1.6 mm (1/16”) thick clear or black cast acrylic (McMaster-Carr, Elmhurst, IL USA) laser-cut with a Trotec Speedy 360 80-Watt laser CO2 engraver/cutter (Marchtrenk, Austria). Parts were custom designed to perfectly fit the InVision stage. For choice and single-cue ethology arenas, three clear pieces were cut: a base with an engraved groove for seating the frame, a frame, and a top with engraved groove and cut inlets for loading agarose casts or parasites. For screening arenas, a single clear base was cut, and a black top was cut with wells designed to fit a multichannel pipette. Each well could fit 16 µL of liquid and avoid creating a meniscus, thus minimizing shadowing in videos. The top was also lightly engraved throughout to aid in adherence during and after fabrication. All arenas limit miracidia movement in the Z-axis, maintaining the parasites within the cameras’ focal plane.

Agarose molds were designed in OnShape (Boston, MA USA) and printed with PLA or PETG filament on Prusa MK3S or XL 3D printers (Prague, Czech Republic). Printers used a 0.4 mm nozzle with 0.2 mm layer height and 15% infill. Molds were designed to create casts that perfectly fit the ethology arenas and hold 180-200 µL of liquid.

Ethology and screening arenas were fabricated by a standard process. Following fabrication, all acrylic pieces were rinsed under tap water to remove residual dust from laser-cutting and then dried thoroughly. For ethology arenas, methyl ethyl ketone (MEK) was applied to the contact surfaces using a micropipette. Approximately 50 µL of MEK was dispensed along each long edge of the frame and 10 µL along each short edge. The frame was aligned with the engraved groove on the base and pressed firmly by hand for several minutes to initiate solvent bonding. Once secure, the top layer was positioned and glued using the same MEK volumes, ensuring proper alignment of the agarose slots and miracidia inlet. Assembled ethology arenas were sealed with tape over the inlets to prevent dust contamination during storage.

For screening arenas, a thin, even layer of MEK was first applied to the bonding surfaces of the clear base and the engraved side of the black acrylic top using a fine paintbrush. Once aligned and pressed together, an additional 100 µL of MEK was slowly pipetted along the entire seam between the two layers to ensure complete bonding. The assembled screening arenas were clamped between two flat wooden boards for several minutes to apply even pressure. Once dry, they were covered with tape and stored in a dust-free container until use.

### Miracidia choice and single-cue ethology assays

Snail-conditioned water (SCW) was prepared by incubating 20 *B. glabrata* NMRI snails (8-15 mm) in 25 mL of APW in a glass beaker for 3 hours at 28°C (10). 2.5 mL aliquots of the resulting solution were lyophilized (Labconco FreeZone, Kansas City, MO USA), and the resulting powder was stored at -80°C.

Agarose casts of cues were made by making 0.5% agarose in APW and pipetting molten agarose into 3D-printed agarose molds. The molds were placed in a humid box and kept in the refrigerator for 12-15 minutes to polymerize. For diffusion tests, 1% Brilliant Blue dye was added to the 0.5% agarose casts. For the choice ethology assays, lyophilized SCW agarose casts were made by adding approximately 180-200 µL of 0.5% agarose solution to a 2.5 mL lyophilized SCW aliquot, mixing the components via pipetting, and adding the mixed solution to the agarose molds. A single 2.5 mL aliquot consists of the combined secretory products from ∼2 snails over 3 hrs. APW agarose casts were also made and used in the choice ethology assays. For the single-cue ethology assays, 2X stocks of each synthesized *Biomphalaria* P12 peptide was made by diluting 2 mg aliquots of the peptides with 1 mL APW. 1 mL of the 2X stocks was mixed with 1 mL of 1% agarose solution, for a final concentration of 1 mg/mL in 0.5% agarose, which was then pipetted into molds.

To prepare for imaging, arenas were wiped down with 70% ethanol and filled with ∼3 mL APW in choice ethology arenas and ∼1 mL APW in single-cue ethology arenas. Agarose casts were added to the filled arenas and incubated at room temperature for 1 hr. to form a cue gradient before adding miracidia. 20 µL of miracidia (>1 hour old) at 1 parasite/µL were added to the entry pores of the arena(s) and placed in InVision for recording. Arenas were transferred to the InVision and recorded for 1 hour at 8 frames per second (FPS). For each treatment condition (raw SCW,

lyophilized SCW with or without pre-diffusion, three independent biological replicates were conducted using miracidia from separate infections.

### High-throughput phenotyping of miracidia

High-throughput behavioral phenotyping was performed with screening arenas and the InVision device. Treatments included four P12 peptides at final concentrations of 200 ng/mL and 400 ng/mL, magnesium chloride at 0.5 mM and 1 mM, and reconstituted lyophilized SCW at volumes of 2 µL and 4 µL per well. Each 10X treatment was prepared in PCR strip tubes, and 1.6 µL of each solution was pipetted into each well of the screening arena using a multichannel pipette (except for SCW, which was pipetted as 2 µL or 4 µL using a single channel pipette). Miracidia were collected after 15 minutes of light induced hatching and suspended in APW at 1 parasite per µL. The miracidia solution was prepared in a 5 mL reservoir with a total volume of 1,500 µL to ensure all 78 wells could be filled. After loading the treatments, 14.4 µL of the miracidia solution was added to each well using a multichannel pipette (12 µL and 14 µL for the 4 µL and 2 µL SCW wells, respectively, to maintain a total volume of 16 µL per well).

For each treatment condition, six technical replicates (wells) were performed per arena, and three independent biological replicates were conducted using miracidia from separate infections. The screening arena was marked in the top left corner to maintain consistent orientation and placed in the InVision system for behavioral recording. Videos were recorded for 5 minutes at 0 (immediately after miracidia were added to pre-loaded wells), 30, 60, 90, and 120 minutes post-treatment. Between recordings, the arena was incubated at 26°C in a sealed chamber with a water reservoir to minimize evaporation.

### Computational tracking and statistical analysis of miracidia in wide-field InVision videos

All analytical scripts and Snakemake workflows are available in the wheelerlab-uwec/invision-tools GitHub repository. Videos from each camera in the 2×2 array were tracked independently and merged afterward if needed. Computational tracking of miracidia is split into two steps, each performed with functions from the soft-matter/trackpy v0.6.4 library (25): segmentation and linking. For segmenting miracidia, the background was generated every 25 frames by taking the maximum projection of a 25-frame chunk. This background was subtracted from each frame in chunk, and tp.batch was used to identify objects in the foreground (diameter = 35, minmass = 1200, noise_size = 2). Object coordinate data was stored in an HDF5 file, which was then amended by the linking step performed by tp.link (search_range = 45, memory = 25, adaptive_stop = 30). The linked objects were stored in a compressed, serialized pickle file which was read by R scripts for all downstream analyses. For videos containing multiple units (single-cue ethology arenas with 3 units per video or screening arenas with 48 units per video), tracks were split with a custom Python script. For ethology arenas that spanned two cameras, tracks from camera pairs were linked to create a single combined pickle file. All tracking workflows were managed by a Snakefile with a modified Slurm executor that sent segmentation jobs to high-memory (>500 GB RAM) nodes and tracking jobs to regular nodes (26,27). Tracking analyses were performed on the BOSE cluster in the Blugold Center for High Performance Computing at UWEC.

Tracking data and analyses can be found in the GitHub repository associated with this manuscript (wheelerlab-uwec/miracidia-sensation-ms) and a Zenodo repository (10.5281/zenodo.15732765). Tracks of miracidia had sporadic gaps resulting from miracidia interactions with the walls of the arena or agarose casts. To account for the wide range of track lengths, only tracks that consisted of >40 frames (∼5 seconds) were used for analyses. Furthermore, each track was split into 5 second chunks and analyzed individually. This chunking scheme ensured that behavioral differences were not artifacts of differences in track length distributions between videos or arena regions and to ensure that long contiguous tracks were properly represented in the final dataset.

Forty-three kinematic, geometric, path, and spatial features were extracted from each 5 second chunk. Each behavioral feature was modeled as a linear mixed effect model (feature ∼ response + (1 + date/video/track) using lme4, lmerTest and the BOBYQ optimizer (28–30) where the fixed response variable was a region (for choice or single-cue ethology arenas) or a treatment (for screening arenas). The model controls for random variation among miracidia extractions and accounts for the nested data structure that includes track chunks that are not independent. Effect sizes were calculated using Cohen’s D, employing the pooled standard deviation for each feature (31). Significant differences in behaviors in ethology arenas were assessed based on a false discovery rate (FDR) threshold of <0.05, and multiple testing in screening arenas was accounted for with the Holm’s correction (32,33).

### P12 cloning, sequencing, and synthesis

Predicted sequences of BGLBO28940 and BGLBO27975 were obtained from the *B. glabrata* BBO2 reference genome (34), available on VectorBase (35), and Primer3 (36) was used to design PCR primers that would amplify the full-length mRNA and the coding sequence of both genes.

RNA was extracted from *B. glabrata* (NMRI), *B. sudanica* KEMRI, and *B. kuhniana* Grande Riviere (37) snails using the Direct-zol RNA Miniprep (Zymo Research, Irvine, CA USA). One *B. glabrata* snail, 1 cm in width, was placed in a glass petri dish, cleaned with 70% ethanol and wiped down using Kimwipes to remove contaminants. *B. sudanica* and *B. kuhniana* snails, obtained from collaborators at the University of New Mexico and stored in Trizol at -80°C, were thawed, removed and cleaned using the same procedure. A small beaker was firmly pressed on top of the snails to shatter the shell, and dissection tools were used to remove the snail tissue from its shell. The tissue was transferred to 1.5 mL RNase/DNase-free tubes and 400 µL of Trizol was added. Sterile plastic pestles were used to homogenize the tissue. The tube was capped, dropped in liquid nitrogen, removed after complete frozen, and homogenized again with the pestle. After homogenization, the tube was centrifuged at 21,300 rcf for 1 min. at room temperature. The supernatant was transferred to a spin column and the manufacturer’s protocol was followed with 10,000 rcf centrifugation steps.

A NanoDrop One (ThermoFisher, Waltham, MA USA) was used to evaluate nucleic acid purity and a Qubit4 fluorometer (ThermoFisher) was used to quantify the extracted RNA. The SuperScript IV First-Strand Synthesis System (ThermoFisher) was used to convert the RNA to cDNA. PCR was used to amplify P12 from the cDNA using the 2X Platinum SuperFi II PCR Master Mix (ThermoFisher). Six reactions were performed in total with the primers designed to amplify the full-length mRNA and the coding sequence for both genes (BGLBO28940 and BGLBO27975) from all three *Biomphalaria* species. Amplification was accomplished using the following cycling conditions: 98°C for 20 seconds, followed by 35 cycles of 98°C for 10 sec., 60°C for 10 sec., 72°C for 30 sec., and finished with 1 cycle at 72°C for 5 minutes and held at 4°C.

Amplified cDNA products were cleaned using the DNA Clean & Concentrator (Zymo). The manufacturer protocol was followed exactly with a 5:1 DNA Binding Buffer to sample ratio. Centrifugation was performed at 10,000 rcf for 30 seconds at room temperature for all steps. Samples were sent to Plasmidsaurus (South San Francisco, CA USA) for amplicon sequencing. A multiple sequencing alignment (MSA) was performed with the sequences from *B. glabrata*, *B. kuhniana*, and *B. sudanica*, and the sequences of *B. straminea* (38) and *B. pfeifferi* (39) using AliView software (40). Four unique versions of P12 were identified from the MSA, and each versions was synthesized for use in ethology assays and penetration experiments (GenScript, Piscataway, New Jersey).

Amplified cDNA products were cloned into pCR-Blunt II-TOPO plasmid vectors using the Zero Blunt TOPO PCR Cloning Kit (ThermoFisher) and transformed into One Shot TOP10 Chemically Competent *E. coli* cells. Steps from the manufacturer’s protocol were followed exactly. The cells were plated on LB + kanamycin (50 µg/mL) agar plates using sterile plating beads and incubated overnight at 37°C. Transformants were picked using sterile pipette tips and placed in 10 mL of LB + kanamycin (50 µg/mL) broth to culture overnight at 37°C in a horizontal test tube shaker set to 180 rpm.

Plasmids of P12 clones were purified using the ZymoPURE Plasmid Prep Kit (Zymo). Steps from the manufacturer protocol were followed exactly. The samples were sent to Plasmidsaurus for sequencing. Sequenced data was uploaded to GenBank (PV848035-PV848039).

### Snail penetration experiments

Wells of a 24-well plate were filled with 1.2 mL of APW, and 5-6 miracidia were individually pipetted into each. After miracidia were added, the peptides were diluted to 1 mg/mL and added to their respective wells for a final concentration of 0.4 mg/mL. An equivalent volume of APW was added to the control well. Lastly, individual *B. glabrata* snails (6-10 mm) were added to each well and the plate was left to incubate at room temperature for 1 hour. After incubation, snails were removed, and the remaining miracidia were stained with Lugol’s solution and counted. Three biological replicates were performed (independent miracidia harvests from different infection cohorts), each with three snails.

## Results

### High-resolution, wide-field imaging of *Schistosoma mansoni* miracidia in custom arenas enables quantitative description of behavioral features

The behavioral and molecular processes involved in schistosome miracidia host-seeking of competent snails has long been of interest but has been limited by technical constraints that inhibit high-resolution tracking of miracidia at relevant spatiotemporal scales. We overcame these constraints through design and fabrication of a custom imaging device combined with bespoke “ethology arenas” produced in-house (Fig 1A-B, S1 Fig). Our InVision (for <u>inv</u>ertebrate <u>vision</u>) is inspired by a similar solution for wide-field simultaneous imaging of *C. elegans* in large phenotypic screens but leverages a unique arena design customized for aquatic organisms (24,41). The InVision deploys a 2×2 array of high-resolution (126.5 px/mm at 20 cm working distance) cameras, utilized in pairs to image two ∼9×3 cm acrylic arenas (Fig 1A). Arenas are laser-cut and adhered with acrylic solvent, with a thin frame separating the top and bottom layers to limit the Z-axis movement of miracidia and ensure consistent localization within the camera’s focal plane. Cut slots in the top layer enable the positioning of agarose casts made with cues of interest. Combination of acrylic arenas and agarose casts produced stable and reproducible gradients (Fig 1B, S1 Movie).

**Fig 1.**
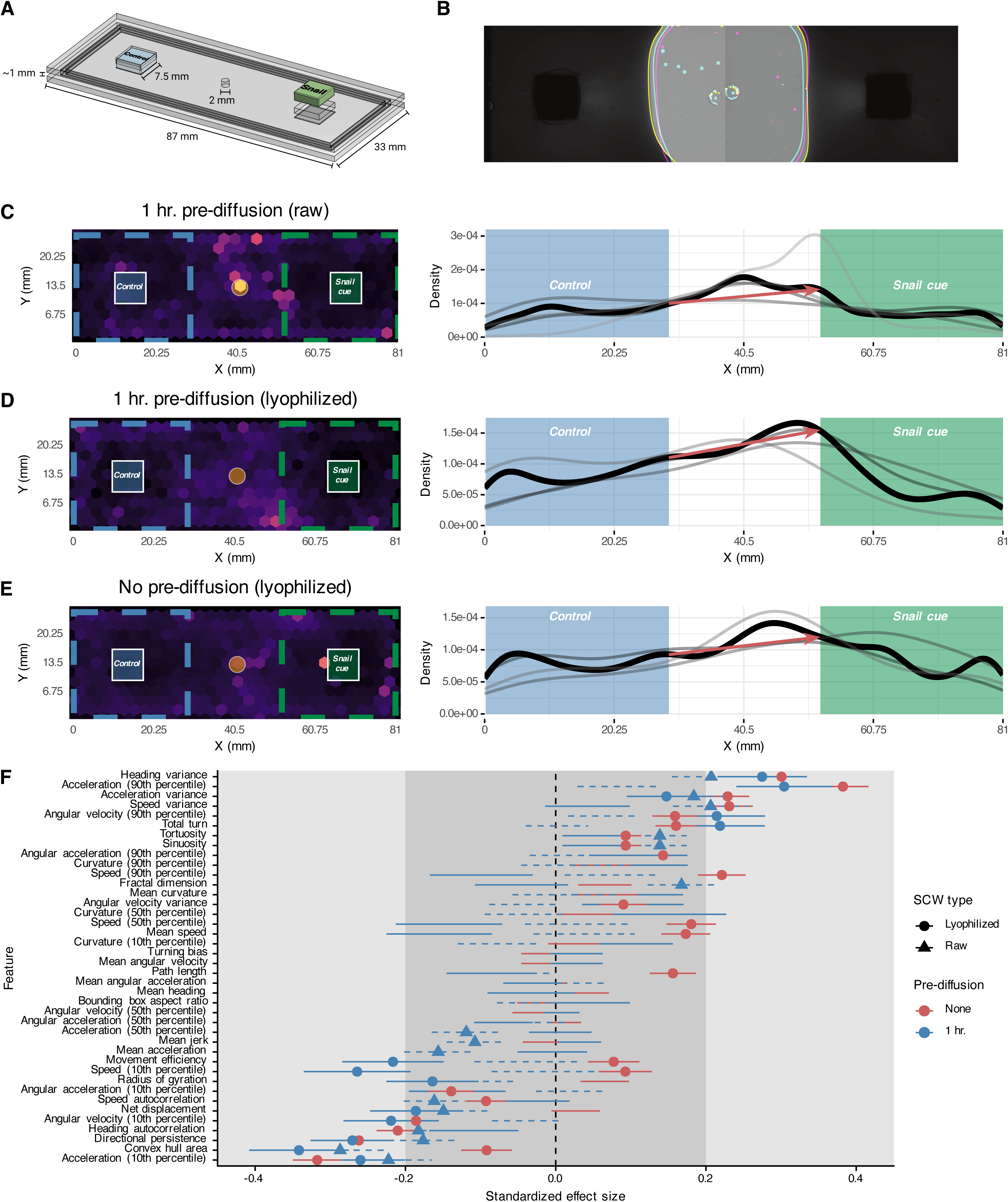
High-resolution tracking of miracidia in a large spatiotemporal scale reveals quantitative ethological features in response to snail cues. (A) Custom choice ethology arena precisely designed to fit the stage of the InVision recording device. Miracidia are loaded through a central pore and allowed to explore the arena. (B) Cue and control gradients generated from an agarose cast in an ethology arena are highly reproducible. Colored lines represent the gradient boundary after 1 hr. diffusion from three independent replicates. (C-E) Left: Heatmaps represent track density over 1 hr. long videos. Right: Density of tracks along the x-axis. Thin grey lines represent independent replicates, and the dark black line represents the mean density across all replicates. Red arrows show the difference in densities at the edge the boundaries. All three panels include data from >3 biological replicates (miracidia cohorts from independent infections). (F) Quantitative features extracted from tracks in the cue or control regions were modeled with linear mixed models. Standardized effect size is Cohen’s D; lines represent the standard deviation of the estimate. Small, medium, and large effect sizes are distinguished by gray boxes. Only lines with points are statistically significant (FDR < 0.05). Positive effect sizes indicate increase in the feature in the cue region when compared to the control region.

To validate our imaging platform, miracidia behavior was recorded in choice ethology arenas using SCW as a cue, which has been shown to cause behavioral changes and accumulation when experienced in a gradient (16,17,42). We used raw, unconcentrated SCW and lyophilized, concentrated SCW to test if SCW concentration influenced miracidia behavior in our experimental setup. We used lyophilized SCW in experiments with and without pre-diffusion of cue. Without pre-diffusion, the concentration gradients were consistently and dynamically expanding (S1 Movie), while with 1 hr. pre-diffusion the gradients stabilized, creating a sharp border between cue, control, and middle regions.

With pre-diffusion of either raw or lyophilized SCW, parasites accumulated along the border of the SCW region, with the starkest difference occurring when using lyophilized SCW, which had the strongest and sharpest gradient (Fig 1C-D). Miracidia also accumulated nearer the SCW region when the cue had not been pre-diffused, but the border accumulation was less pronounced (Fig 1E). Interestingly, in no experiment did miracidia accumulate next to the cue source, which is consistent with the steep gradient supported by our arenas and the “boundary reaction” described in classical ethological studies, which maintains miracidia in the active space of a snail (16,43).

A key advance is our wide-field tracking approach is the ability to extract quantitative behavioral features that describe the kinematics, path geometries, and spatial and angular dynamics of miracidia tracks. Because we track miracidia in the entire arena and maintain stable, non-overlapping gradients, we can compare behavioral features and assess chemoklinokinesis in different regions of interest. We computed a set of 43 quantitative features from short track segments (“chunks”) and modeled each feature using a linear mixed-effects model to test for regional behavioral differences while accounting for within-video and within-individual variation. After correcting for multiple comparisons, we found many behavioral features that were significantly different between cue and control regions for experiments with pre-diffusion (lyo: 14 features, raw: 16 features, FDR < 0.05) and without pre-diffusion (25 features) (Fig 1F). In all experiments, heading variance and acceleration (90^th^ percentile) had the greatest positive effects, while directional persistence, convex hull area, and acceleration (10^th^ percentile) had the greatest negative effect. Overall, the behavioral profiles were remarkably similar between experiments, with only two of the features showing opposite effects. Interestingly, where significantly different features showed the same directional effect in all experiments, the miracidia in arenas without pre-diffusion routinely exhibited greater changes in behavior (as measured through absolute value of standardized effect size), which may be explained by the dynamic gradient requiring a more refined and sensitive response (Fig 1F).

In contrast to the few quantitative behavioral features that have been analyzed in the past, accumulation at a gradient has been a consistent observation in descriptive studies of miracidia behavior. However, how this accumulation is accomplished has not been made clear. Some reports suggest that moving up the gradient does not lead to a behavioral change; instead, moving down the gradient will lead to behavioral changes that result in accumulation and maintenance within the active space of the cue (16,44). Others describe behavioral changes when moving in either direction (42,45). Our choice ethology arenas allow clear comparisons between the behaviors exhibited while moving in or out of a cue region. When comparing only track chunks that crossed a cue or control boundary in pre-diffused experiments (739 total chunks), we found more behavioral features that were significantly different for tracks moving out of a cue region (16 features) than into a cue region (11 features) (Fig 2A). Remarkably, while the overall profile was similar for the two behaviors, there were six features that were affected in the opposite direction and were greatly dissimilar – miracidia moving out of a region (“in to out”) had more turns and a higher variance in the angular velocity (Fig 2B). These differences support the previously described “boundary reaction,” which ensures that miracidia do not leave an active space after serendipitously discovering it. Our data suggests that miracidia perform small to moderate changes in behavior when experiencing either environmental change – moving into or out of a cue – but will substantially increase their turning when leaving a cue.

**Fig 2.**
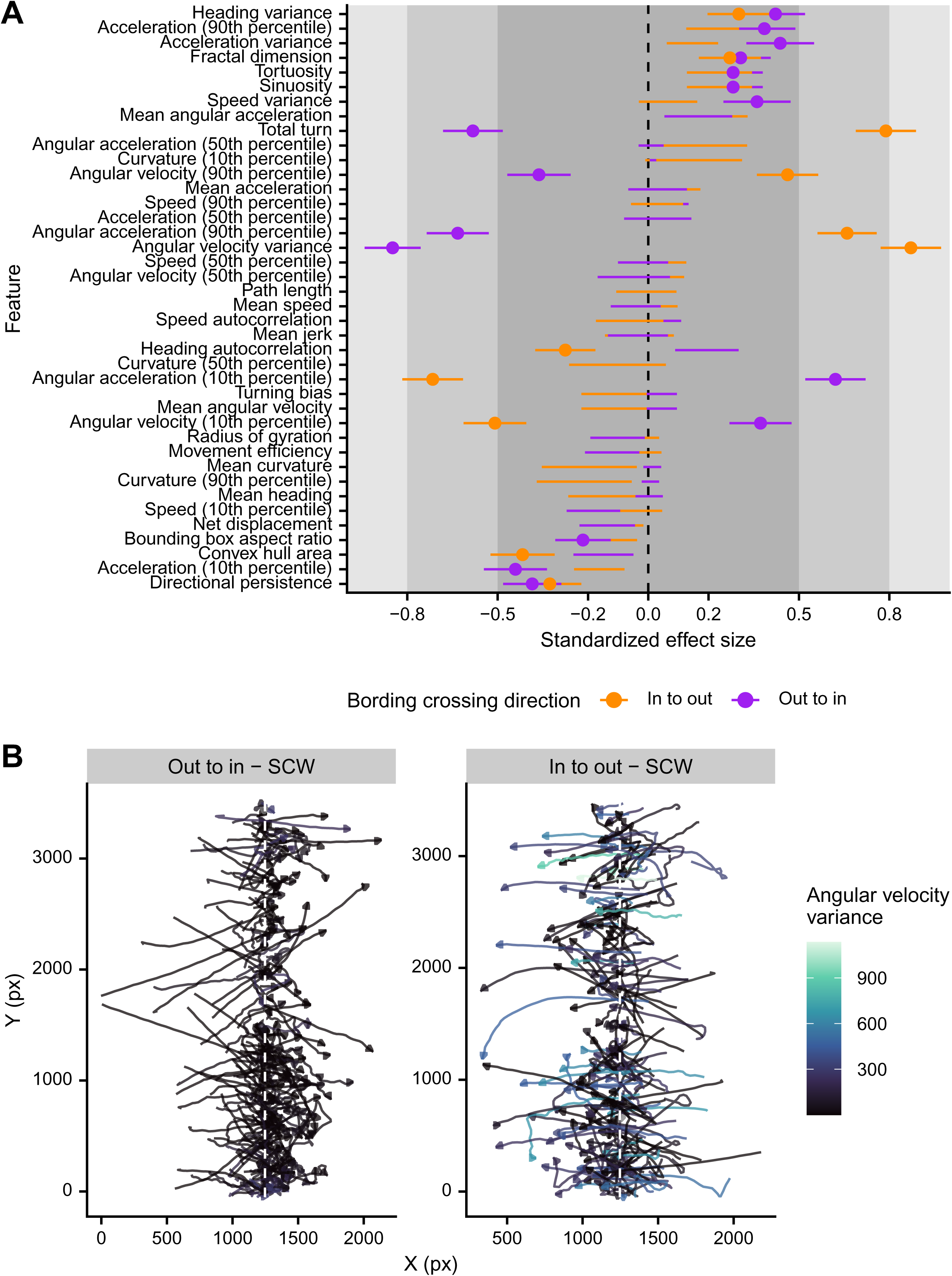
Quantitative description of the border response of miracidia moving in or out of a cue region. (A) Quantitative features were extracted from track chunks crossing the border of cue or control regions and modeled as a response of region. Standardized effect size is Cohen’s D; lines represent the standard deviation of the estimate. Small, medium, and large effect sizes are distinguished by gray boxes. Only lines with points are statistically significant (FDR < 0.05). Positive effect sizes indicate increase in the feature when crossing the cue boundary when compared to the control boundary. (B) Extracted border crossing track chunks showing the increased turning of miracidia moving out of the cue region in comparison to those moving into the region.

### Natural variation in a conserved *Biomphalaria* peptide causes divergent behavior and accumulation effects on *S. mansoni* miracidia

Our novel platform and initial experiments with SCW began to unravel the quantitative ethology of the miracidia response to snail cues, and we next moved to exploring the role of specific snail cues that may be involved in eliciting the miracidia response. Much effort has been given to identifying molecules within SCW that contribute to the behavioral effect, with data suggesting that glycoproteins, salts, or even protons could be involved (9,46–48). Recently, a secreted snail peptide (P12) was identified in fractionated *B. glabrata* SCW and shown to cause accumulation and behavioral change when miracidia were exposed to it in a gradient (10,11,21). To allow side-by-side comparisons of multiple cue sources rather than a binary choice between two cues, we designed single-cue ethology arenas to simultaneously record miracidia from the same cohort (Fig 3A). Like the choice ethology arenas, we verified that stable and reproducible gradients could be produced (S2 Movie).

**Fig 3.**
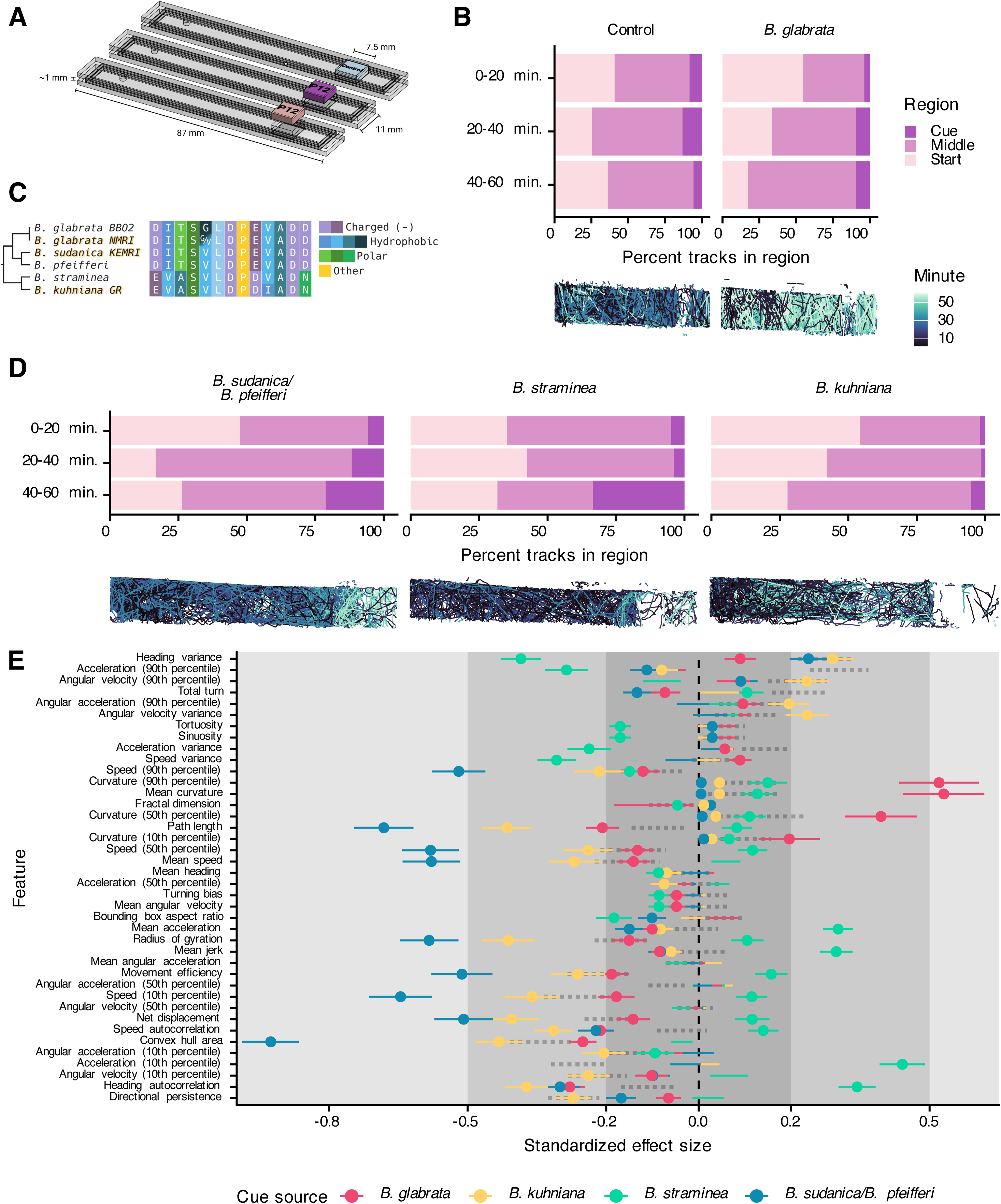
Single-cue ethology arenas allow side-by-side comparisons of the quantitative behavioral features of *Biomphalaria* spp. P12 peptides. (A) Custom single-cue ethology arenas allow for simultaneous imaging of three arenas per InVision camera pair. (B) Top: Number of tracks per region (1/3 of the arena) over time. Bottom: Representative tracks. Miracidia progressively accumulate nearer the BgP12 cue but not the control cue. (C) Multiple sequence alignment of cloned and reference P12 homologs. Gold-shadowed sequences were cloned. Cladogram derived from (49). (D) Same as (B) but with new *Biomphalaria* homologs. (E) Extracted behavioral features from track chunks in the cue region were compared to control experiments. Standardized effect size is Cohen’s D; lines represent the standard deviation of the estimate. Small, medium, and large effect sizes are distinguished by gray boxes. Only lines with points are statistically significant (FDR < 0.05). Positive effect sizes indicate increase in the feature in the cue region when compared to the control region. Features on the y-axis are in the same order as Fig 1F; dotted lines represent the values from lyophilized SCW in Fig 1F.

We initially tested these arenas by comparing the response of miracidia to a gradient of BgP12. When compared to the control, miracidia progressively accumulated in regions nearer the BgP12 source over time (Fig. 3B). Behavioral features of chemoklinokinetic responses were significantly altered when miracidia were in the presence of BgP12. Although the behavioral profile was similar to that caused by lyophilized SCW, several features were markedly different (Fig. 3E).

At the time of discovery, BgP12 was not known to have homologs in other snails due to a lack of gastropod genomic resources. Using the *B. glabrata* BB02 reference sequence for P12 (34), we found homologs in *B. pfeifferi* (39), *B. straminea* (38), and *B. sudanica* 111 (50) reference genomes. Notably, we did not find a P12 homolog in the *Bulinus truncatus* genome (51), which is the primary African vector for *Schistosoma haematobium* and a planorbid related to *Biomphalaria*, or the genome of *Oncomelania hupensis*, the vector of *Schistosoma japonicum* and not closely related to the planorbid schistosome vectors (52). Thus, P12 seems to be restricted to *Biomphalaria*. Using primers designed using the full-length mRNA that encodes the putative BgP12 pro-peptide, we cloned sequences from *B. glabrata* NMRI, *B. sudanica* KEMRI, and *B. kuhniana* GR (37) (GenBank accession numbers: PV848035- PV848039). Amongst these five related *Biomphalaria* snail species, four versions of the P12 peptide were identified and varied at seven residues, as the P12 from the closely related *B. pfeifferi* and *B. sudanica* were identical. Our lab strain of *B. glabrata* NMRI was found to be heterozygous at one position in the P12 peptide sequence (Fig 3C).

We synthesized these peptides and used them in single-cue ethology assays. The *B. sudanica/B. pfeifferi* and *B. straminea* sequences caused clear accumulation over the length of the experiment, while the *B. kuhniana* peptide had a moderate effect similar to the control (Fig 3D). These results suggest that accumulation is not a general response to a general peptide gradient but is specific to the P12 sequences. These data partially correlate with what is known about compatibility between *S. mansoni* and *Biomphalaria* spp. *S. mansoni* NMRI readily infects *B. glabrata, B. pfeifferi, B. sudanica,* and *B. straminea* but does not infect *B. kuhniana* (37).

We extracted features from tracks and compared differences between tracks on the cue-half or entry-half of the arenas. There was not a generic behavioral profile in response to P12s, and profiles varied from the lyophilized SCW profile (Fig 3E). Instead, the *B. sudanica/B. pfeifferi* and *B. straminea* peptides, which caused the greatest amount of accumulation near the cue, caused very different behavioral features but consistently resulted in higher effect sizes than the *B. glabrata* or *B. kuhniana* homologs. Even though *B. kuhniana* did not cause strong accumulation, it still caused some behavioral change, but most features differences were of small effects (Fig 3E).

These data indicated that all four *Biomphalaria* P12 variants elicit chemoklinokinetic behavioral features but different behavioral profiles. Miracidia accumulation nearer the cue region also differed, with the *B. sudanica/B. pfeifferi* P12 causing the most accumulation while also leading to the greatest effects on chemoklinokinetic features. Since accumulation is an emergent property of chemoklinokinetic features, it is notable that the cues that caused the greatest accumulation also had largest effects on quantitative features.

### Snail-derived cues only cause miracidia chemoklinokinesis when presented in a gradient

To combat schistosomiasis in regions with persistent hotspots, the World Health Organization recommends snail control through molluscicide treatment or habitat removal (53). An alternative to widespread extermination of an important ecological regulator may involve reduction in the number of infected snails rather than reduction of the snail population as a whole (54). Inhibition of miracidia chemoklinokinetic features and resulting accumulation in snail active spaces may achieve this result, either through direct blocking of the chemoklinokinetic process elicited by snail cues, synthetic stimulation of the chemoklinokinetic process to stop miracidia dispersal and entry into the snail active space, or through masking of natural snail gradients. Based on data showing that P12 homologs cause accumulation and diverse behavioral effects, we reasoned that P12 may be a potential chemoklinokinetic stimulant.

To determine an effective concentration for P12 exposure, we designed a high-throughput behavioral assay that would enable dose-response experiments. Our custom screening arena maintains the advances of the ethology arenas (no meniscus, limited Z-axis movement) while allowing 96-wells to be simultaneously recorded by each camera pair (Fig 4A). Tracking miracidia over 5 minute videos results in high-quality, contiguous paths (Fig 4B).

**Fig 4.**
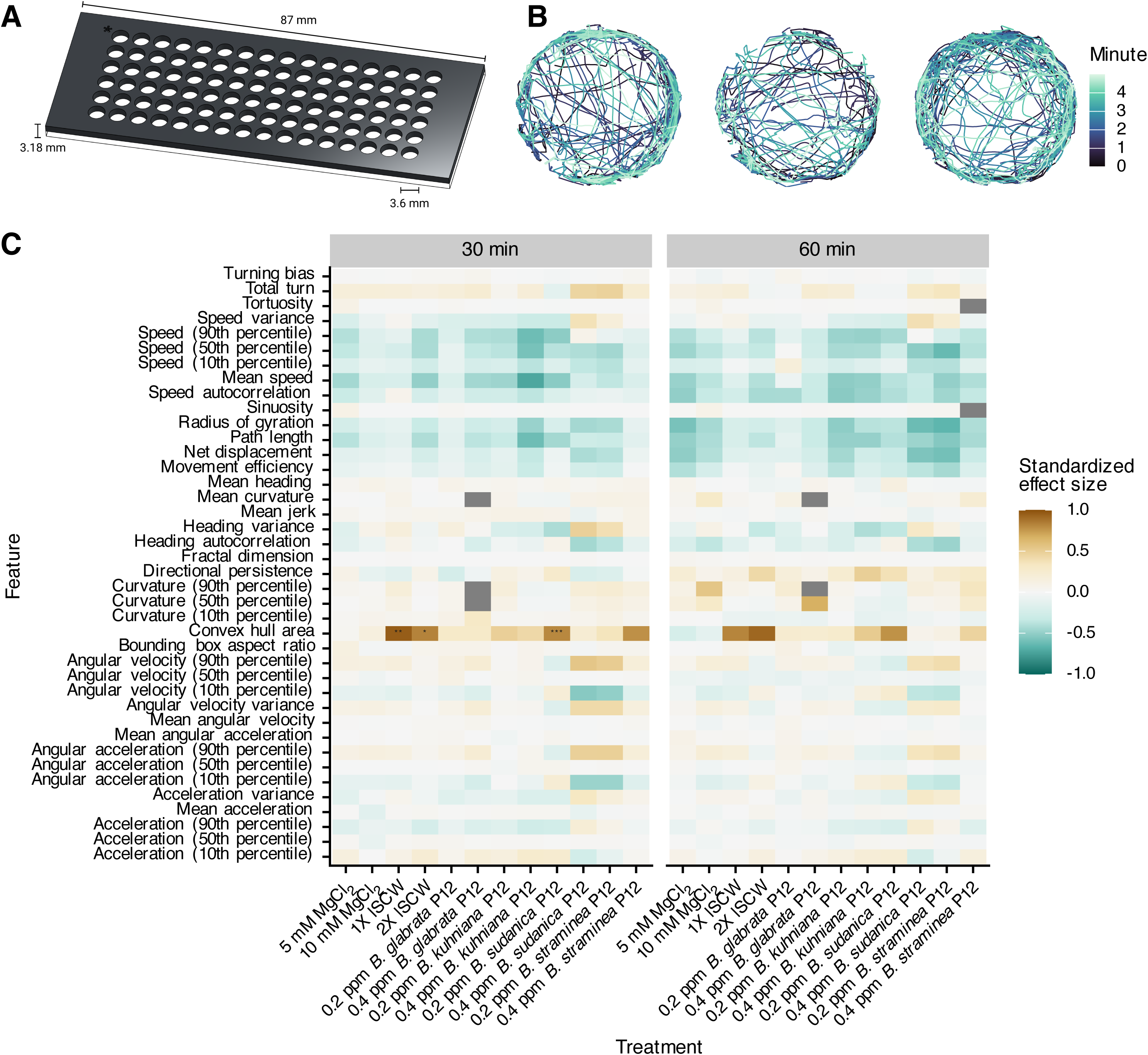
High-throughput behavioral screening of chemoklinokinetic stimulants. (A) Custom screening arena for simultaneous recording of 96 wells per InVision camera pair. (B) Representative tracks of three wells. (C) Extracted behavioral features from track chunks in the treated wells were compared to control wells. Standardized effect size is Cohen’s D. Positive effect sizes indicate increase in the feature in the treated wells when compared to the control wells. P-values are adjusted with Holm’s correction (*p ≤ 0.05; **p ≤ 0.01, ***p ≤ 0.001).

We exposed miracidia to P12 at concentrations consistent with those experienced in ethology arenas. As potential positive controls, we also used lyophilized SCW and MgCl2, which has previously been shown to drive chemoklinokinetic behavior and accumulation (18,55). We recorded miracidia immediately after exposure and every 30 minutes for 90 minutes. After 30 or 60 minutes, most quantitative features were not significantly different from control wells (Fig 4C).

There were some differences immediately after exposure, but the effect sizes were small and were lost by 30 minutes (S2 Fig). We reproduced these results with SCW and MgCl2 using droplets on a slide to ensure that negative results were not an artifact of our screening arena(56) (S3 Fig). These data show that the behavioral changes exhibited in ethology arenas are dependent upon a gradient – uniform presentation of the cue does not cause chemoklinokinesis. The changes caused by SCW and P12 peptides are thus truly sensory in nature and not a generic neuromuscular stimulation, as the gradient requirement has been repeatedly shown in other metazoan systems that use chemotaxis (57–59). Notably, the contrast in miracidia behavior when in a gradient or in a uniform concentration has been a topic of some disagreement in the literature (14).

### Uniform presentation of miracidia to *Biomphalaria* P12s inhibits penetration of snails

The inability to activate chemoklinokinesis in high-throughput context impeded our ability to identify effective concentrations that would act as chemoklinokinetic stimulant to block dispersal in a natural environment. However, these results led us to reason that P12 peptides may be able to mask the natural snail cue gradient and inhibit the behavioral processes that lead to penetration and infection. Miracidia exhibit chemoklinokinetic responses when they enter the active space created by natural snail secretions. In this theoretical framework, a masked gradient would cause miracidia to exhibit normal exploratory behavior instead of chemoklinokinesis, even after having entered a snail active space (Fig. 5A).

**Fig 5.**
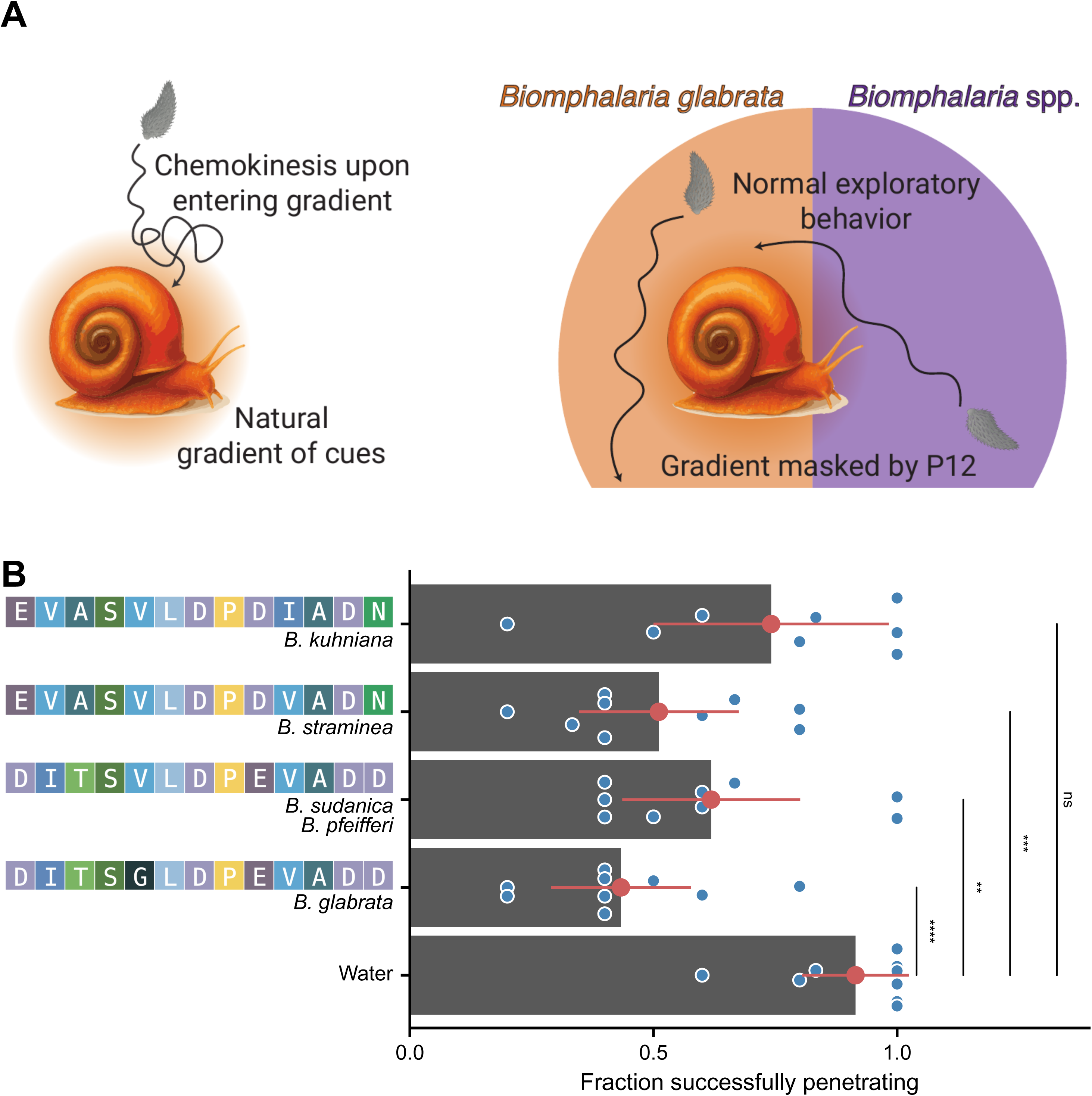
Inhibition of snail penetration by gradient masking. (A) Theoretical model for how *Biomphalaria* P12s may block miracidia penetration of snails. (B) Fraction of miracidia remaining in well after 1 hr. incubation with snails in the presence or absence of *Biomphalaria* P12 peptides. Each point represents an independent penetration experiment (snail in a well with miracidia). Data is the combination of three biological replicates (three snails exposed to miracidia from independent infection cohorts). Points and ranges represent the mean and confidence intervals. P-values are the result of t-tests (*p ≤ 0.05; **p ≤ 0.01, ***p ≤ 0.001; ****p ≤ 0.0001).

To test our hypothesis, we conducted penetration experiments with miracidia and snails in the presence or absence of P12 peptides. After incubating for 1 hour, 90% of untreated miracidia penetrated the snail tissue. When we added *Biomphalaria* P12s, the peptides from *B. glabrata, B. straminea,* and *B. sudanica/B. pfeifferi* significantly decreased the number of miracidia that penetrated the snail after 1 hour (Fig 5B). BgP12 was the most effective, causing a >50% reduction in penetration. Remarkably, *B. kuhniana* P12, which differs from *B. straminea* by only a single amino acid and *B. glabrata* by 5 amino acids, did not have a significant effect on miracidia penetration (Fig 5B). These data are consistent with the ethology experiments, which showed that BkP12 caused less accumulation than the other P12s and had smaller effects on quantitative behavioral features (Fig 3).

## Discussion

We developed a scalable imaging and analytical platform that enables high-resolution, quantitative ethology of schistosome miracidia. This approach captures their behavior at previously inaccessible spatiotemporal scales and helps reconcile decades of fragmented observations in behavioral parasitology.

Beyond basic discovery, these capabilities support translational research by enabling the identification and characterization of compounds that either stimulate or block chemoklinokinetic responses. Either approach may be able to effectively block snail infection, which would be a major advance and additional tool for schistosomiasis control. Given the ecological importance of aquatic snails, our strategy for reducing the number of infected snails rather than the general snail population may offer a preferable alternative for control approaches targeting the snail stages. Quantitative ethology of miracidia in response to prioritized compounds may help predict effective infection blockers.

There remains opportunity for further advancements. Our finding that snail cues fail to induce chemoklinokinesis under uniform presentation poses a challenge for high-throughput screening. Effective inhibitor screening requires a consistent, reproducible activation of behavior, which is currently limited by the need for stable gradients. It remains unclear whether this limitation applies only to sensory-level stimuli or also to downstream neuromodulators. Compounds that act beyond the initial sensory detection may still prove effective. To overcome this, future screens could leverage microfluidic technologies that support miniaturized gradient formation.

While our study identifies behavioral signatures linked to successful penetration and further implicates conserved *Biomphalaria* P12 peptides as potential modulators of host-finding, their molecular targets in miracidia remain unknown. Uncovering how miracidia detect these cues could employ integration of single-cell transcriptomics, peptide-receptor mining, and new functional tools (60–62). Although reverse genetics in larval schistosomes remains technically challenging, our platform offers a robust foundation for bridging behavior with molecular mechanisms in this host-parasite interface.

## Acknowledgements

*Schistosoma mansoni*-infected mouse tissue was provided by the Schistosomiasis Resource Center of the Biomedical Research Institute (Rockville, MD) through NIH NIAID Contract HHSN272201700014I. Funding was provided by Student Blugold Commitment Differential Tuition funds through the University of Wisconsin-Eau Claire and the UWEC Office of Academic Affairs. Some computational resources used for this study were provided by the Blugold Center for High-Performance Computing under NSF grant CNS-1920220. NJW, RVH, and CNN were supported by NIH NIAID R15 AI183095. The authors would like to thank Dr. Jennifer Smith (University of Wisconsin-Eau Claire) for use of her freeze dryer, Drs. Martina Laidemitt and Sam Loker (University of New Mexico) for frozen samples of *Biomphalaria sudanica* and *Biomphalaria kuhniana* samples, Max Hofbauer (loopbio) for InVision design services, and the UWEC Makerspace for laser-cutting and some 3D printing services.

