## Supplementary figures and images for "Quantitative ethology of schistosome miracidia characterizes a conserved snail peptide that inhibits penetration"

### S1 Movie

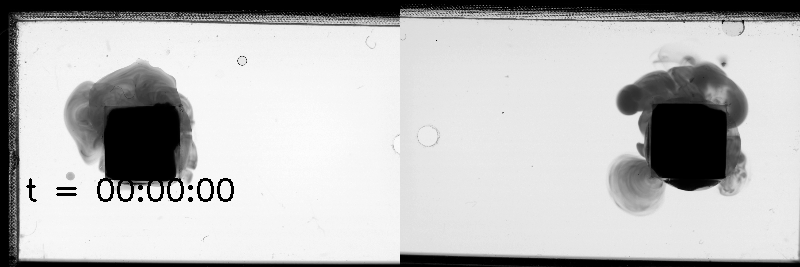

### S2 Figure

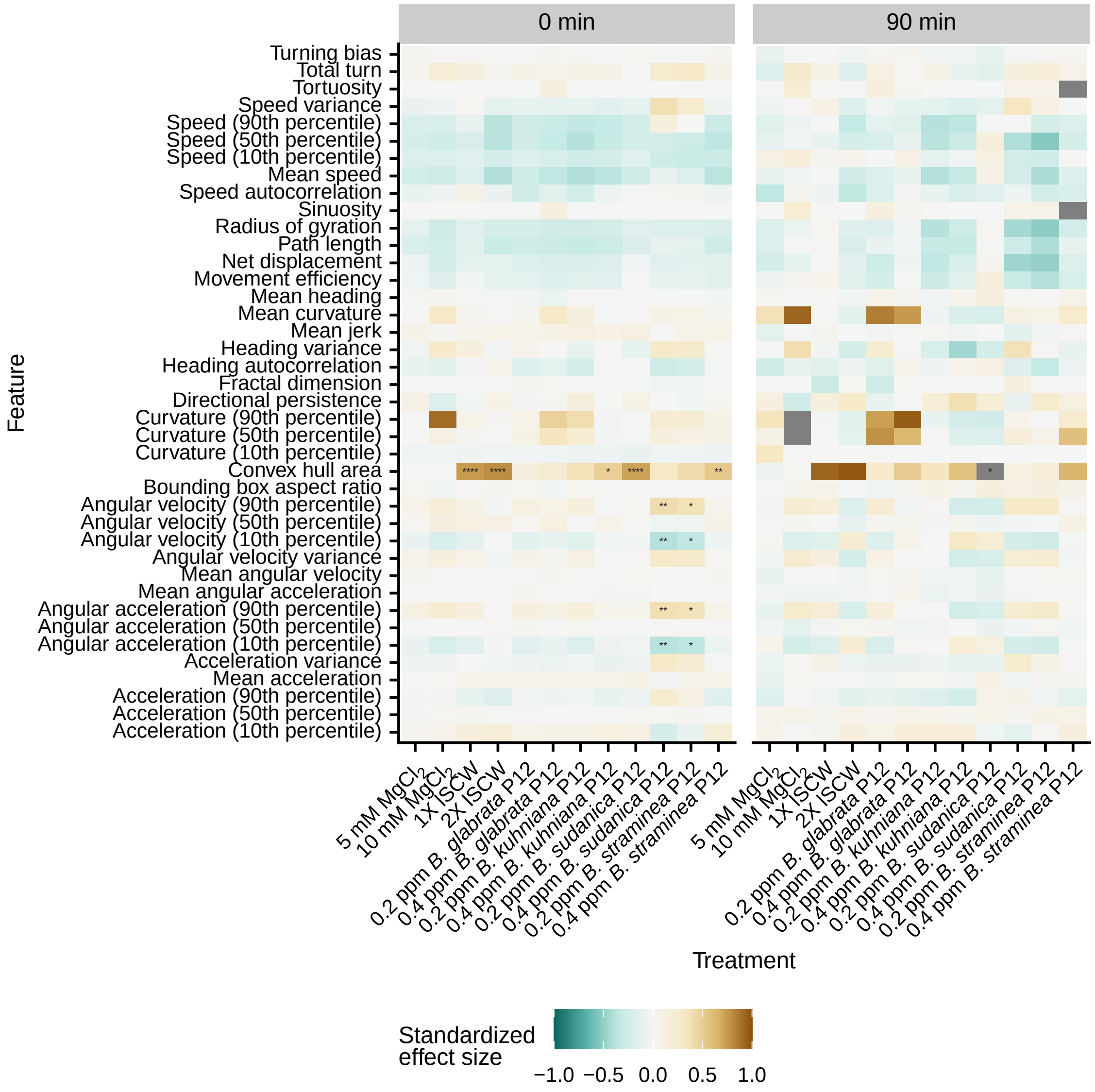

### S2 Movie

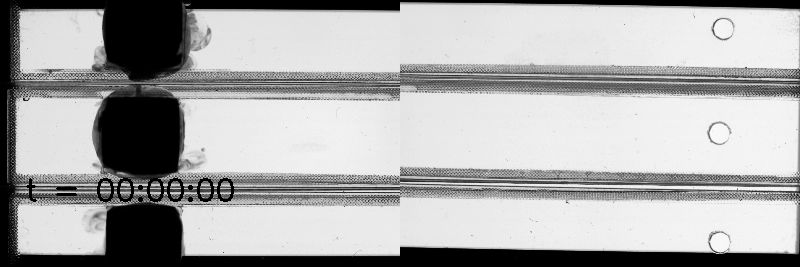

### S3 Figure

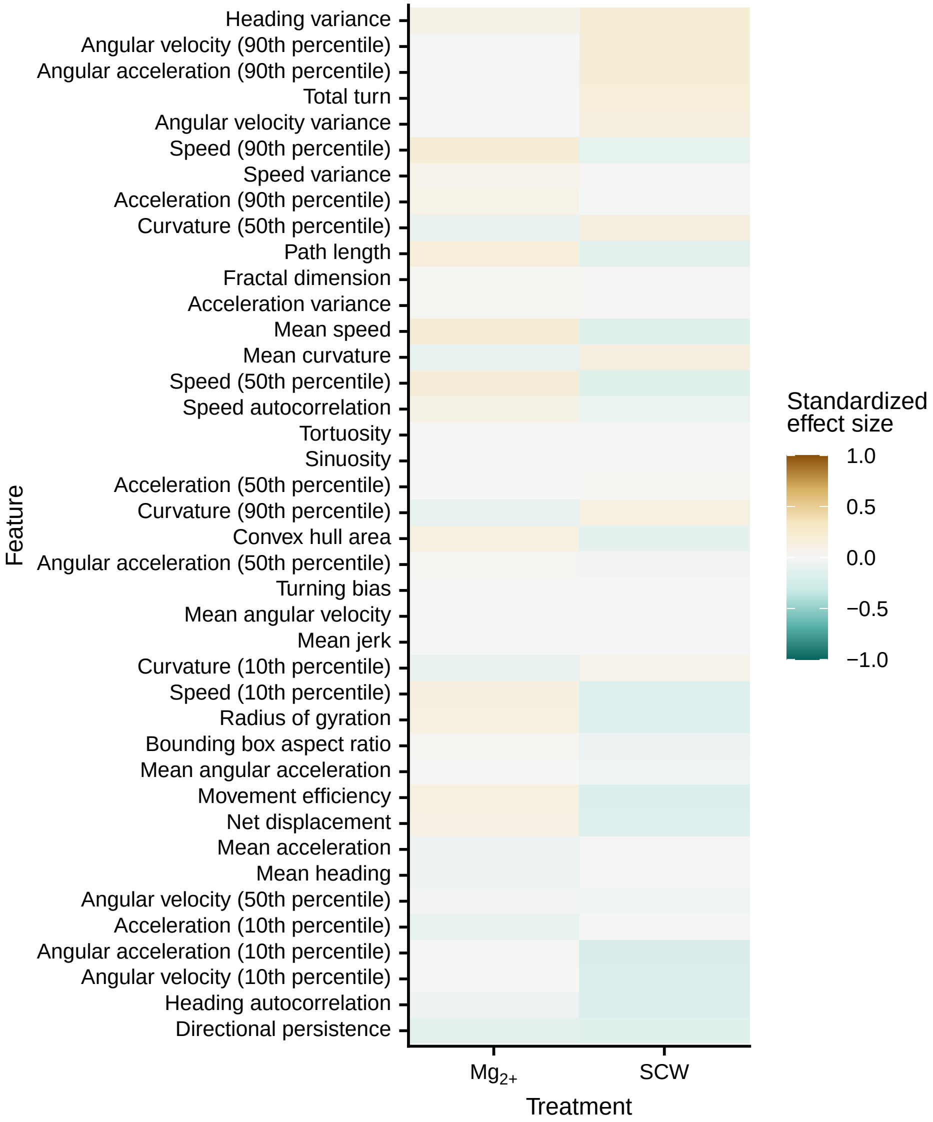
